# Rapid evolution of bacterial AB5 toxin B subunits independent of A subunits: sialic acid binding preferences correlate with host range and toxicity

**DOI:** 10.1101/2021.05.28.446168

**Authors:** Naazneen Khan, Aniruddha Sasmal, Zahra Khedri, Patrick Secrest, Andrea Verhagen, Saurabh Srivastava, Nissi Varki, Xi Chen, Hai Yu, Travis Beddoe, Adrienne W. Paton, James C. Paton, Ajit Varki

**Affiliations:** Glycobiology Research and Training Center, University of California San Diego, San Diego, California, USA; Department of Cellular & Molecular Medicine, University of California San Diego, San Diego, California, USA; Department of Chemistry, University of California Davis, Davis, California, USA; Department of Biochemistry and Molecular Biology, Monash University, Clayton, VIC, 3800, Australia; Department of Animal, Plant and Soil Science and Centre for Agri Bioscience (Agri Bio), La Trobe University, Bundoora, VIC, 3086, Australia; Research Centre for Infectious Diseases, Department of Molecular and Biomedical Science, University of Adelaide, Adelaide, SA 5005, Australia

**Keywords:** Evolution, Bacterial, Toxin, Phylogenetic, *Yersinia pestis*, Host Range, Sialoglycan Microarray, Cytotoxicity, Pathogenesis

## Abstract

Cytotoxic A subunits of bacterial AB_5_ toxins enter the cytosol following B subunit binding to host cell glycans. We report that A subunit phylogeny evolves independently of B subunits and suggest a future B subunit nomenclature based on species name. Phylogenetic analysis of B subunits that bind sialic acids (Sias) with homologous molecules in databases also show poor correlation with phylogeny. These data indicate ongoing lateral gene transfers between species, with mixing of A and B subunits. Some B subunits are not even associated with A subunits e.g., YpeB of *Yersinia pestis*, the etiologic agent of plague epidemics. Plague cannot be eradicated because of *Y. pestis*’ adaptability to numerous hosts. YpeB shares 58% identity/79% similarity with the homo-pentameric B subunit of *E. coli* Subtilase cytotoxin, and 48% identity/68% similarity with the B subunit of *S*. Typhi typhoid toxin. We previously showed selective binding of B_5_ pentamers to a sialoglycan microarray, with Sia preferences corresponding to hosts e.g., *N*-acetylneuraminic acid (Neu5Ac; prominent in humans) or *N*-glycolylneuraminic acid (Neu5Gc; prominent in ruminant mammals and rodents). Consistent with much broader host range of *Y. pestis*, YpeB binds all mammalian sialic acid types, except for 4-O-acetylated ones. Notably, YpeB alone causes dose-dependent cytotoxicity, abolished by a mutation (Y77F) eliminating Sia recognition, suggesting cell proliferation and death *via* lectin-like cross-linking of cell surface sialoglycoconjugates. These findings help explain the host range of *Y. pestis* and could be important for pathogenesis. Overall, our data indicate ongoing rapid evolution of both host Sias and pathogen toxin-binding properties.

## Introduction

Secreted bacterial AB_5_ toxins are an important class of virulence factors, named after their unique architecture, typically comprising a single catalytically active toxic A subunit and pentameric B subunits (1–3). Such AB_5_ toxins classically exert effects on host cells via two steps: the five B subunits form a ring-shaped pentamer responsible for binding host cell glycans and mediating uptake of holotoxin; this is followed by inhibition of host cellular functions mediated by toxic effects of the A subunit. Several of these toxins are known to recognize specific sialylated glycan moieties on target cells through a glycan-binding pocket in their B subunit (4–7). The notable exception is Shiga toxin, which binds globoseries glycolipids (1).

AB_5_ toxins are typically classified and named based on the catalytic activity of their A subunits. The cholera toxin family includes Ctx and *Escherichia coli* heat-labile enterotoxins (LTs) LT-I and LT-II (8), and a less-well characterized toxin from *Campylobacter jejuni*. These toxins all have A subunits with ADP-ribosylation activities targeting an arginine of G_s_α (the α subunit of the stimulatory trimeric G protein) (1). Pertussis toxin (Ptx) produced by *Bordetella pertussis* (the causative agent of whooping cough) has a catalytic subunit with similar ADP-ribosylase activity, but is placed in a separate family because in contrast to all the other AB_5_ toxins, its B subunit is hetero-rather than homo-pentameric (9). A further AB_5_ toxin variant (referred to as Plt [Pertussis like toxin] or typhoid toxin) is produced by the typhoid fever-causing agent *Salmonella enterica* serovar Typhi (*S*. Typhi). Plt has an A subunit (PltA) with ADP-ribosylase activity and a homopentameric B subunit (PltB), which collectively act as a delivery vehicle for an additional toxic subunit related to cytolethal distending toxin, resulting in a novel A_2_B_5_ architecture (6). The Shiga toxin (Stx) family, produced by *Shigella dysenteriae* and certain strains of *E. coli*, have A subunits with RNA-*N*-glycosidase activity that inhibits eukaryotic protein synthesis (1). A subset of Shiga toxigenic *E. coli* (STEC) strains also produce subtilase cytotoxin (SubAB) the prototype of a fourth AB_5_ toxin family. The A subunit SubA is a subtilase family serine protease with exquisite specificity for the essential endoplasmic reticulum chaperone BiP/GRP78 (3, 10). An additional family is exemplified by an extra-intestinal *E. coli* toxin EcxAB, the A subunit of which is a metzincin-type metalloprotease (11). Microbes that express AB_5_ toxins have a wide range of pathological effects on human populations, from travelers’ diarrhea caused by enterotoxigenic *E. coli* strains to the more serious and life-threatening diarrhea caused by *Vibrio cholerae* (12, 13), and systemic disease such as typhoid fever and the hemolytic uremic syndrome triggered by distinct *E. coli* strains producing Stx and possibly also SubAB (5, 14).

Despite sharing a similar structural architecture, the B pentamers of the various AB_5_ family members differ widely in their host cell surface receptor specificity and intracellular trafficking. For example, PltB selectively recognizes the sialic acid *N*-acetylneuraminic acid (Neu5Ac), which is very prominent in humans because of an inactivating mutation in the *CMAH* gene encoding the CMP-*N*-acetylneuraminic acid hydroxylase that converts Neu5Ac to *N*-glycolyl-neuraminic acid (Neu5Gc) in most other mammals (15). On the other hand, *E. coli* SubB prefers binding to Neu5Gc over Neu5Ac (5). SubB is closely related to PltB (50% identity, 68% similarity over 117 amino acids). However, SubB is even more closely related (56% identity, 79% similarity over 136 aa) to a putative exported protein of *Yersinia pestis* (Accession No. YPO0337, now WP_002209112) (10), although curiously, there is no associated A subunit.

*Yersinia pestis*, the etiologic agent of plague was responsible for major devastating epidemics in human history that claimed millions of lives (16) and is still considered of global importance to public health and potentially in biowarfare. It was first isolated by Alexandre Yersin during the third pandemic plague that reached Hong Kong (17). Plague is predominantly a zoonosis and a *Y. pestis* reservoir is maintained in animals such as ground squirrels and prairie dogs and transmitted to humans by fleas. *Y. pestis* also circulates in its rodent reservoir by flea-borne transmission (18). One very interesting feature of *Y. pestis* is its adaptability to more than 200 hosts (19). In addition to *Y. pestis*, the genus *Yersinia* also includes *Y. pseudotuberculosis* and *Y. enterocolitica*, which are also associated with human infections and cause mild diarrhea (20). *Y. pestis* is thought to be a “young” pathogen that diverged from the more environmental stress-tolerant and less pathogenic bacterium *Yersinia pseudotuberculosis* around 5,000 to 7,000 years ago by adapting to a flea-transmitted life cycle and the capability to cause host systemic infection (21, 22). *Y. pestis* is now considered a clonally expanded genomically degenerating variant of *Y. pseudotuberculosis*. During its evolution, *Y. pestis* showed a varied mutation rate, not strictly obeying a constant evolutionary clock (23). Three forms of plague are usually described, including bubonic, pneumonic, and septicemic plague (20). *Y. pestis* can be transmitted by flea bites causing lymph nodes to swell and form buboes (known as bubonic plague) and respiratory droplets from one person to another (causing pneumonic plague). However, when *Y. pestis* spreads via blood it causes septicemic plague (20). Humans get infected either by flea bites or contact with the infected pets/domestic animals (causing conjunctivitis, skin plague, or pneumonic plague) (24). If bubonic plague is not recognized and treated in time, it can develop into pneumonic plague or systemic plague. Septicemic plague has a very high mortality rate and can also be caused directly by blood infection with the pathogen through a cut (25).

We report here for the first time that the putative *Y. pestis* B subunit binds to and is intrinsically toxic towards cells that express Neu5Ac and/or Neu5Gc. These findings are consistent with *Y. pestis* adaptability of having more than 200 hosts and provide insights into the molecular bases for its host specificity, in understanding fundamental cellular functions and potential treatment of the disease.

## Results and Discussion

### A and B subunits of AB_5_ bacterial toxins evolve independently of each other

With the development of high throughput technologies and well-organized databases, there is a major increase in availability of bacterial AB_5_ toxin sequence data. Based on phylogenetic analysis of existing genomes, it is evident from Fig. 1A that there is a lack of evolutionary relationship between the A and B subunits. For example, while the A subunit (ArtA) of *S*. Typhimurium ArtAB shares its clade with its homologue in *S. bongori*, the B subunit (ArtB) shares a clade with a homologue from *Y. enterocolitica* and *E. coli* EcPltB (Fig. 1A, also detailed in the companion paper, Sasmal et al., 2020). Likewise, while the *E. coli* toxin SubAB shares little similarity with other species with respect to its A subunit (SubA), it shares a clade with proteins from Yersinia species with respect to the B subunit (SubB). This situation has likely arisen because of lateral transmission and mixing and matching of subunit-encoding genes during the ongoing evolutionary arms-race of bacterial species and toxins with respect to the host specificity. The fact that many AB_5_ toxin operons are encoded by bacteriophages and/or conjugative plasmids means that their distribution among species is also influenced by plasmid compatibility and phage host specificity.

**Fig. 1.**
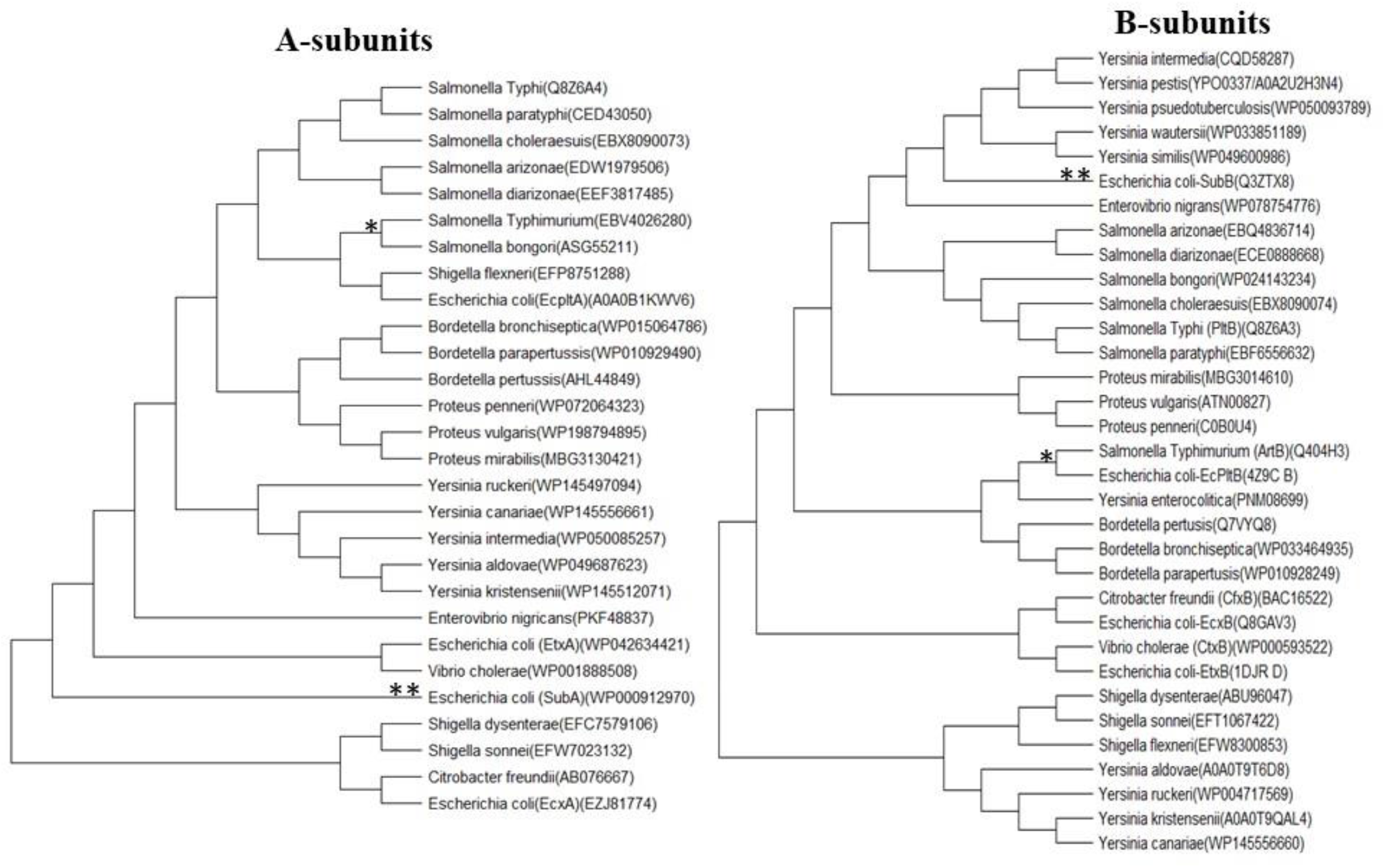
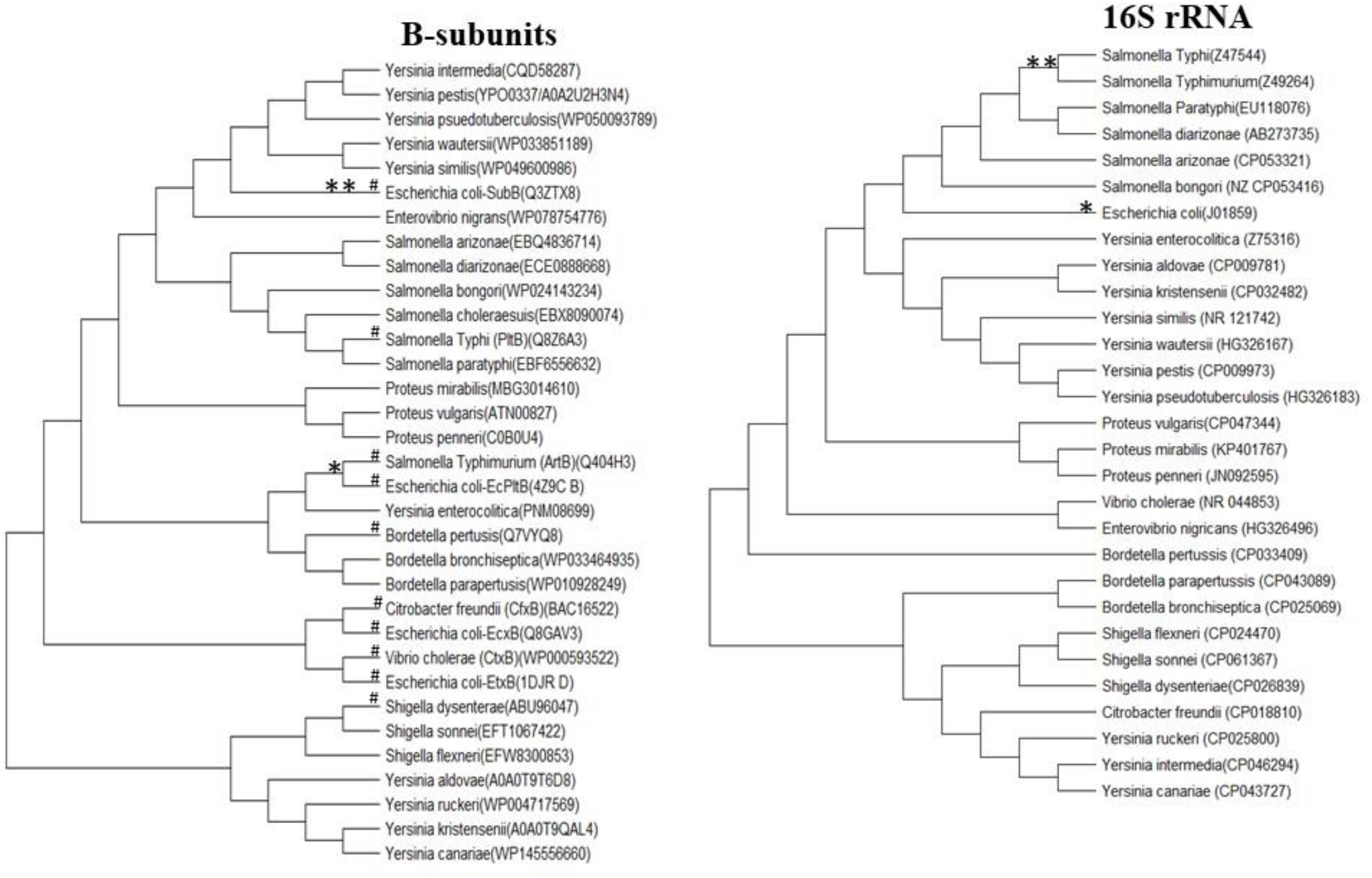
**A. Comparative phylogenies of A and B subunits of AB_5_ bacterial toxin show independent evolution of both subunits.** * shows phylogenetic relationship of *S*. Typhimurium with other bacterial species with respect to A and B subunit and **indicates *E. coli* sharing its clade with other bacterial species based on A and B subunit. (see details of the methodology in “experimental procedures”). **B. Phylogenies of AB_5_ toxin B subunit showing partial discordance with the species phylogeny based on 16S rRNA.** * shows *S*. Typhimurium phylogenetic relationship with other bacterial species with respect to B subunit and 16S rRNA and **indicates *E. coli* sharing its clade with other bacterial species based on B subunit and 16S rRNA. (see details of the methodology in “experimental procedures”). # represents published AB_5_ toxin B subunits with known host Sialic acid binding patterns.

### Phylogenies of B subunits are also partially discordant with species phylogeny

To understand the phylogeny of the various B subunits, we further compared it with species phylogeny based on 16S rRNA. It is evident from the phylogenetic tree (Fig. 1B) that B subunits are also evolving relatively independent of each other, in relation to species phylogeny. From the phylogenetic tree (Fig. 1B), there is a partial discordance even between B subunit and species phylogeny. For example*, E. coli* SubB shares a clade with homologues produced by *Yersinia* species, whereas in species phylogeny, the *E. coli* strains that produce SubB share a clade with *Salmonella* species. Similarly, *S*. Typhimurium ArtB shares its clade with homologues in *Y. enterocolitica* but shares similarity with other Salmonella species based on 16S rRNA. Since the A and B subunits are independent and their biological roles are also distinct, a B subunit nomenclature based on the A subunit seems no longer logical. On the other hand, making changes in existing terminology will lead to confusion regarding the prior literature. We suggest that existing names remain unchanged going forward, but that the nomenclature of an entirely novel toxin should be related wherever practical to the bacterial species producing it (see Table 1). Accordingly, the novel exported protein that appears to be an isolated *Y. pestis* B subunit is hereby named YpeB (see Table 1). Having said that, the lab of one of the co-authors previously referred to *S*. Typhi PltB as *S*. Typhi ArtB in literature (26), because of the close similarity of both the A and B subunits to ArtAB of *S*. Typhimurium (already described by Saito et al., 2005) and the clear dissimilarity between the *S*. Typhi <u>homo</u>pentameric B subunit and the <u>hetero</u>pentameric B subunit of pertussis toxin (Plt stands for pertussis-like toxin).

**Table 1.** Current and Proposed Nomenclature of B_5_ Subunit Toxins

| Bacterial Species | Holotoxin Name | A subunit name | B Subunit name | Citation |
| --- | --- | --- | --- | --- |
| *Vibrio cholerae* | Cholera Toxin | CtxA | CtxB | (50) |
| *Salmonella enterica* serovar Typhi | Typhoid Toxin | PltA | PltB<br>PltC | (6, 51) |
| *Yersinia pestis* |  | - | YpeB | (10) |
| *Yersinia enterocolitica* |  | - | YenB | This study |
| *Salmonella Typhimurium* |  | ArtA | ArtB | (7) |
| Enterotoxigenic *Escherichia coli* | Labile enterotoxin I<br>Labile enterotoxin II | LT1A<br>LT2A | LT1B<br>LT2B | (52) |
| Shiga toxigenic *E. coli* | Shiga toxin 1<br>Shiga toxin 2<br>Subtilase cytotoxin | Stx1A<br>Stx2A<br>SubA | Stx1B<br>Stx2B<br>SubB | (10, 53) |
| Extra-intestinal *E. coli* |  | EcxA<br>EcPltA | EcxB<br>EcPltB | (11, 54) |

### YpeB exhibits broad specificity for sialic acid terminated glycans

Genomic database searches carried out as part of the original characterization of SubAB (10) identified the presence of the closely related proteins encoded by *Y. pestis* (YPO0337; 56% identity, 79% similarity over 136 aa) and *S*. Typhi (STY1891; 50% identity, 68% similarity over 117 aa), now referred to as YpeB and PltB, respectively. Interestingly, it was noted that unlike all other AB_5_ toxins known at the time, the gene encoding YpeB was not closely associated with an A subunit gene (10). This observation raises questions about its role (if any) in pathogenesis of the disease and whether it binds to host sialoglycans. PltB has been reported to bind to the human-dominant sialic acid (Neu5Ac) but not to the other major mammalian sialic acid Neu5Gc (4). Conversely, *E. coli* SubB strongly prefers to bind to the non-human Neu5Gc over Neu5Ac (5). In order to determine the glycan binding patterns of YpeB, we used a customized sialoglycan array (27) to compare the ability of His_6_-tagged-YpeB and fluorescently labeled anti-His secondary antibody to bind pairs of sialylated glycans terminating in either Neu5Ac (prominently expressed in human cells) or Neu5Gc (prominently expressed in cells of most other mammals). Consistent with *Y. pestis* having >200 hosts (rats, rabbits, squirrels, chipmunks etc.), YpeB bound to both Neu5Ac and Neu5Gc terminated glycans (detailed in Fig. 2) implying a potentially broad range of host specificity. Notably, this toxin did not bind to 4-O-acetyl-Neu5Ac (Neu4,5Ac2) or -Neu5Gc (Neu5Gc4Ac).

**Fig. 2.**
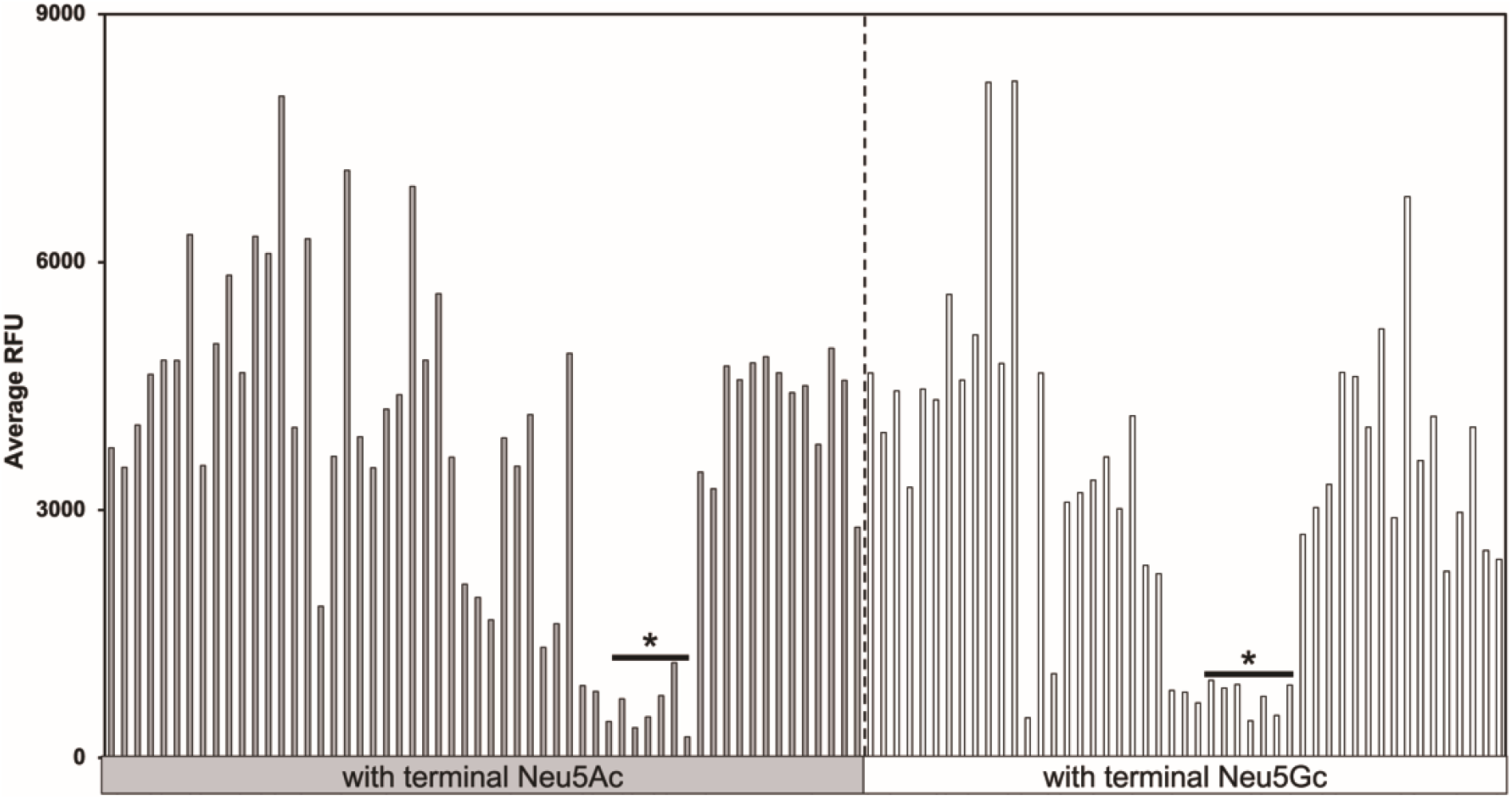
YpeB exhibits strong binding to all Neu5Ac- and Neu5Gc-terminated glycans (except 4-O acetylated, shown by asterix *). The right-side bar shows Neu5Gc-terminated glycans and left side bar shows Neu5Ac-terminated glycans.

### B subunit clustering based on sialic acid binding pattern

With the three already published toxins (PltB, SubB and ArtB) having diverse sialoglycan binding preferences and host range specificity (4–7), we decided to compare the sialoglycan binding preference of these toxins with that of YpeB (Fig. 3). We confirmed previous findings that PltB binds to the human-dominant sialic acid Neu5Ac but not to the other major mammalian sialic acid Neu5Gc, explaining a potential role for Typhoid toxin in pathogenesis in humans (humans get typhoid fever, but our close evolutionary relatives do not) (4). *E. coli* strains producing SubAB are commonly found in ruminants and have been associated with severe human disease (28). However, these strains also produce one or more members of the Shiga toxin family (StxAB), and so although purified SubAB is highly lethal for laboratory mammals (13), the importance of SubAB to pathogenesis of human disease is uncertain. Interestingly, however, this toxin prefers to bind to non-human Neu5Gc over Neu5Ac (5). The AB_5_ toxin ArtAB (29) is produced by *S*. Typhimurium, which is found in a broad range of species from cattle to humans and is a common cause of gastroenteritis. Its B subunit (ArtB) binds to both Neu5Ac and Neu5Gc (7). Interestingly, YpeB also binds to Neu5Ac and Neu5Gc and has an even broader range of host specificity. Detailed comparative sialoglycan binding of these toxins is shown in the following companion paper (Sasmal et al., 2021).

**Fig. 3.**
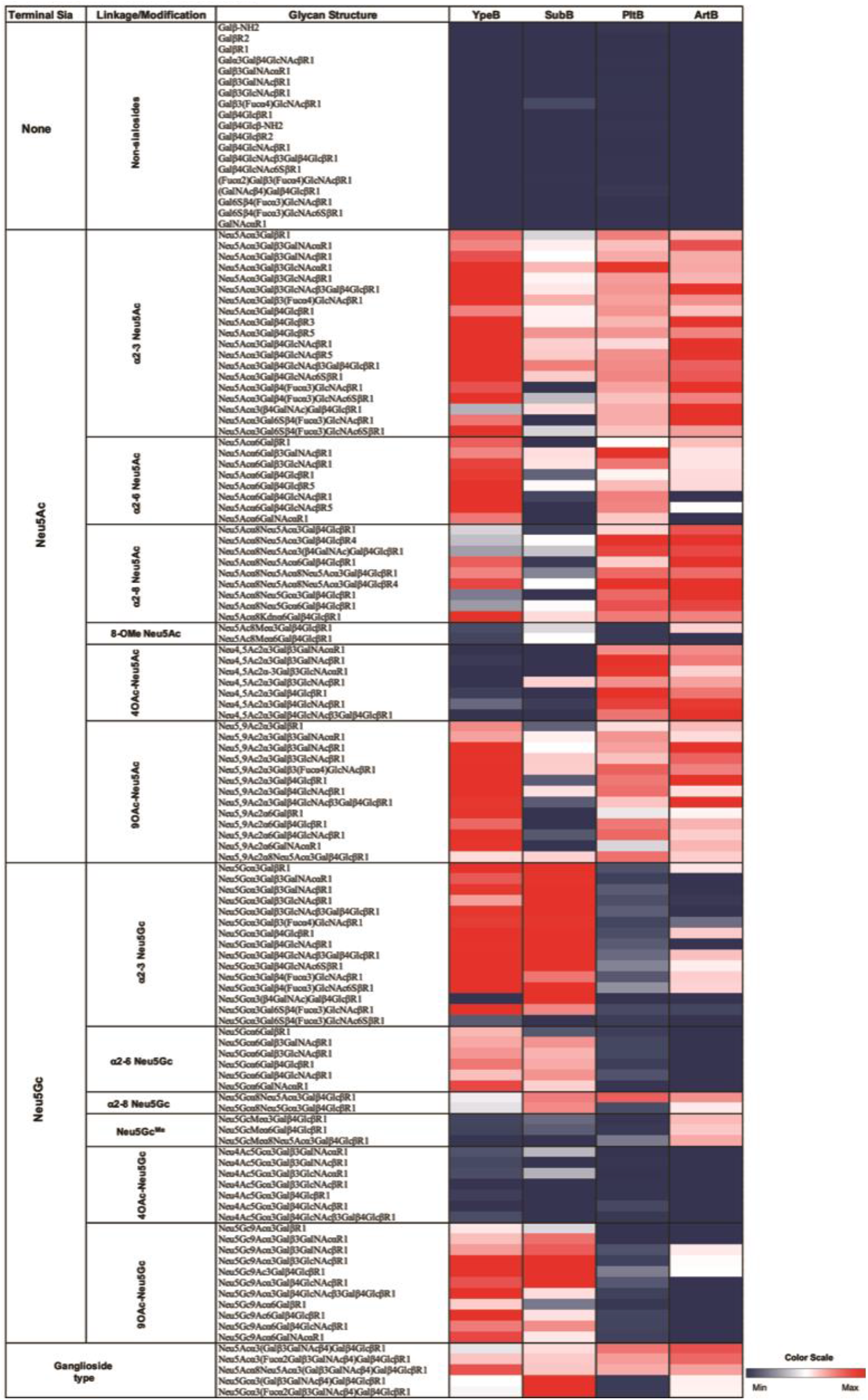
Comparative toxin clustering of PltB, SubB, ArtB and YpeB based on their sialic acid binding pattern. Average relative fluorescence unit (RFU) analyzed from Glycan array are shown. The heatmap show RFU value ranges from lowest (Black) to highest (Red).

### Sialic acid dependent binding of YpeB in CHO cells

To understand if the sialoglycan binding pattern exhibited by YpeB is Sia-dependent, we used hyposialylated LEC29.Lec32 CHO cells (30) and fed them with exogenous Neu5Ac or Neu5Gc and then exposed them to different concentrations of YpeB. As evident from Fig. 4, the binding of YpeB to cells increases with increasing dosage of YpeB when fed with 3 mM Neu5Ac (Fig. 4). Moreover, a higher dosage of YpeB (100 μg/ml) is required to bind to Neu5Gc-fed cells as compared to Neu5Ac-fed cells (Fig. 4).

**Fig. 4.**
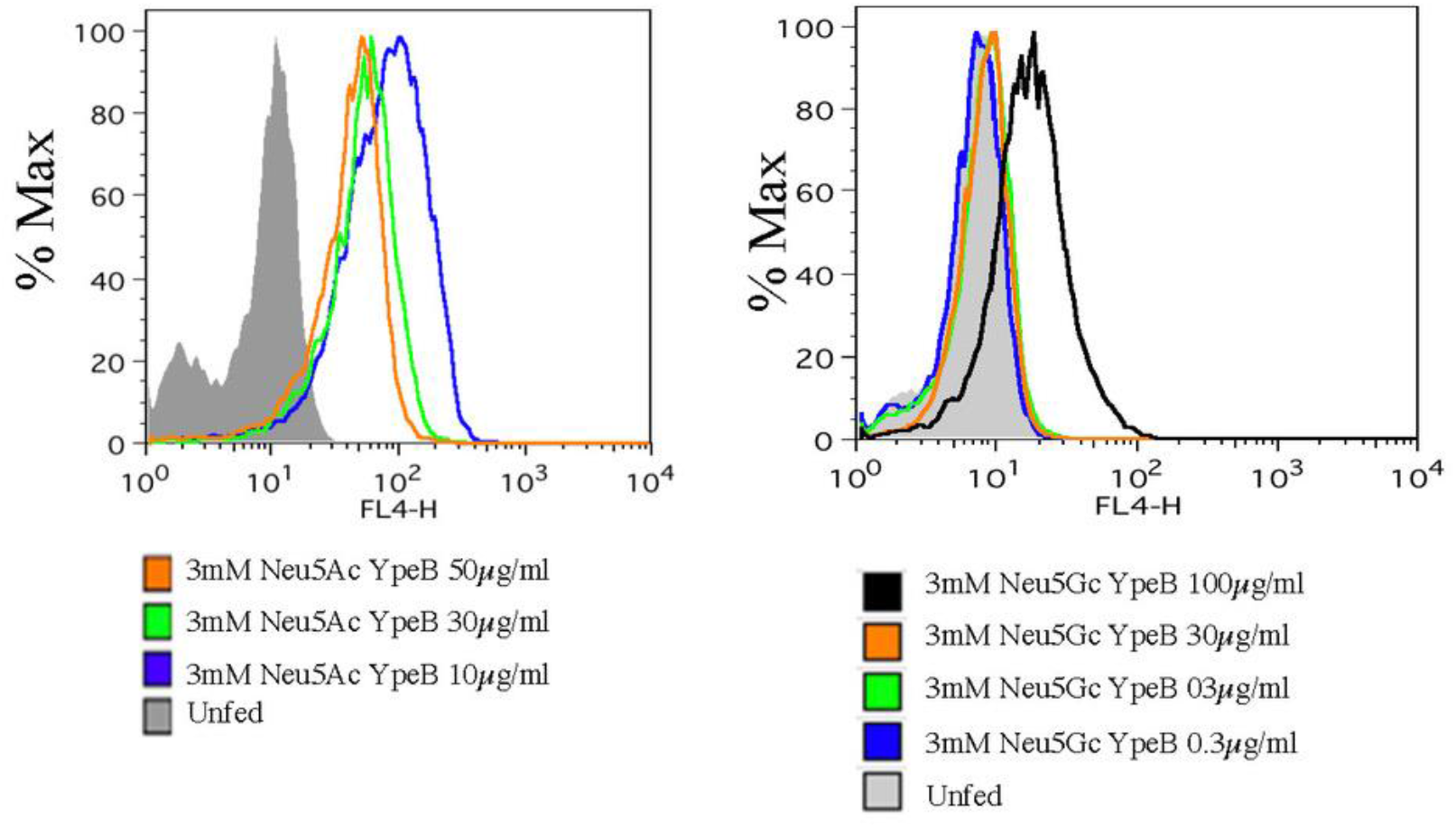
Sialic acid dependent binding of YpeB toxin with hyposialylated LEC29.Lec32 CHO cells fed with Neu5Ac. The cells were cultured with 3mM Neu5Ac for 16hr probed with different concentrations of YpeB (10,30,50 μg/ml). Similarly, the cells were cultured with 3mM Neu5Gc for 16hr probed with different concentration of YpeB (0.3,3,30,100 μg/ml) and binding was analyzed by flow cytometry.

### Structural modelling and YpeB mutant impact on glycan binding

Since the amino acid sequence of YpeB shares 56% identity and 79% similarity with SubB, we predicted the structure feature and amino acids of YpeB that are crucial for binding to host glycans using SubB as a template in Chimera (Fig. 5A). It is well documented that two amino acids (S12 and Y78) are crucial for the binding of glycans to SubB, with a S12A substitution abolishing glycan binding completely, while a Y78F mutation selectively abolished binding to Neu5Gc (5). Taking advantage of structural modelling, we predicted that Y77 in YpeB might also be critical for binding to host glycans. Indeed, when we mutated the amino acid residue in this position from tyrosine (Y) to phenylalanine (F) (Y77F), binding of glycans was abolished, explaining the essential role of Y77 in binding to host cell surface glycans (Fig. 5B). Comparative structural modelling of published toxin B subunits (PltB, SubB, ArtB, YpeB) reported in Fig. 3 are shown in Fig. 6. It is clear from Fig. 6 that PltB lacks an essential tyrosine that could be the reason for its selective binding to human dominant Neu5Ac. Unlike PltB, SubB contains both essential serine and tyrosine residues in its glycan binding pocket, with the latter required for its capacity to preferentially bind Neu5Gc over Neu5Ac (5). Similarly, the ArtB structural model shows both serine and tyrosine in the glycan binding pocket, and in addition also contains an extra loop that could contribute to its broad glycan target specificity.

**Fig. 5A.**
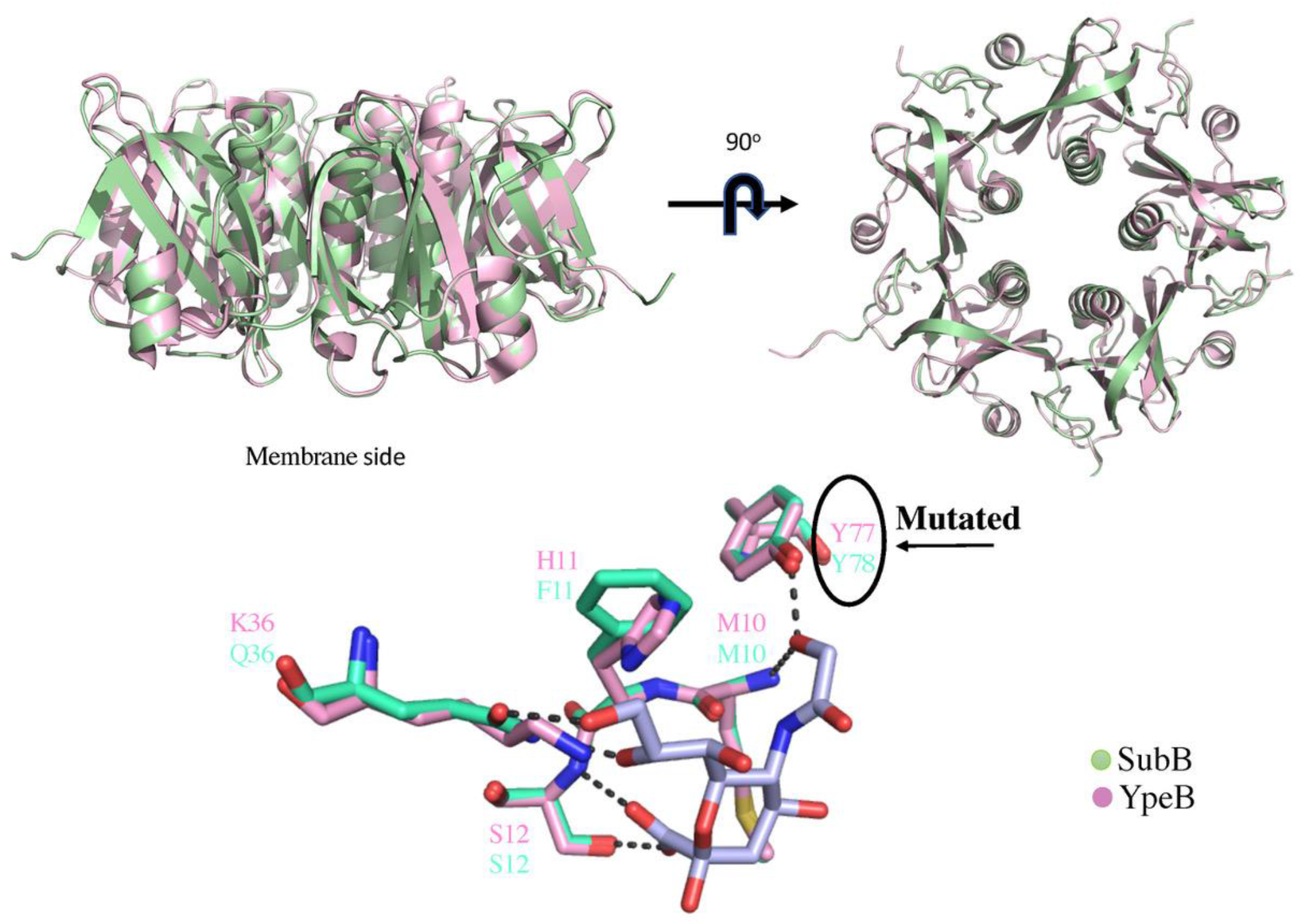
Structural modelling of YpeB toxin based on *E.coli* SubB (PDB ID: 4DWA) as a template. The sequences of *E. coli* SubB and novel YpeB is aligned, and structural modelling was done superposing SubB X-ray structure (pale pink) (PDB ID:3DWP) to the model of YpeB (pale green) using the C-alpha coordinates of all residues. Sialic acid binding pocket indicates Tyrosine 77 (marked as black arrow) to be essential for binding to host glycans. Pink color depicts *Yersinia pestis* YpeB and green shows *E. coli* SubB.

**Fig. 5B.**
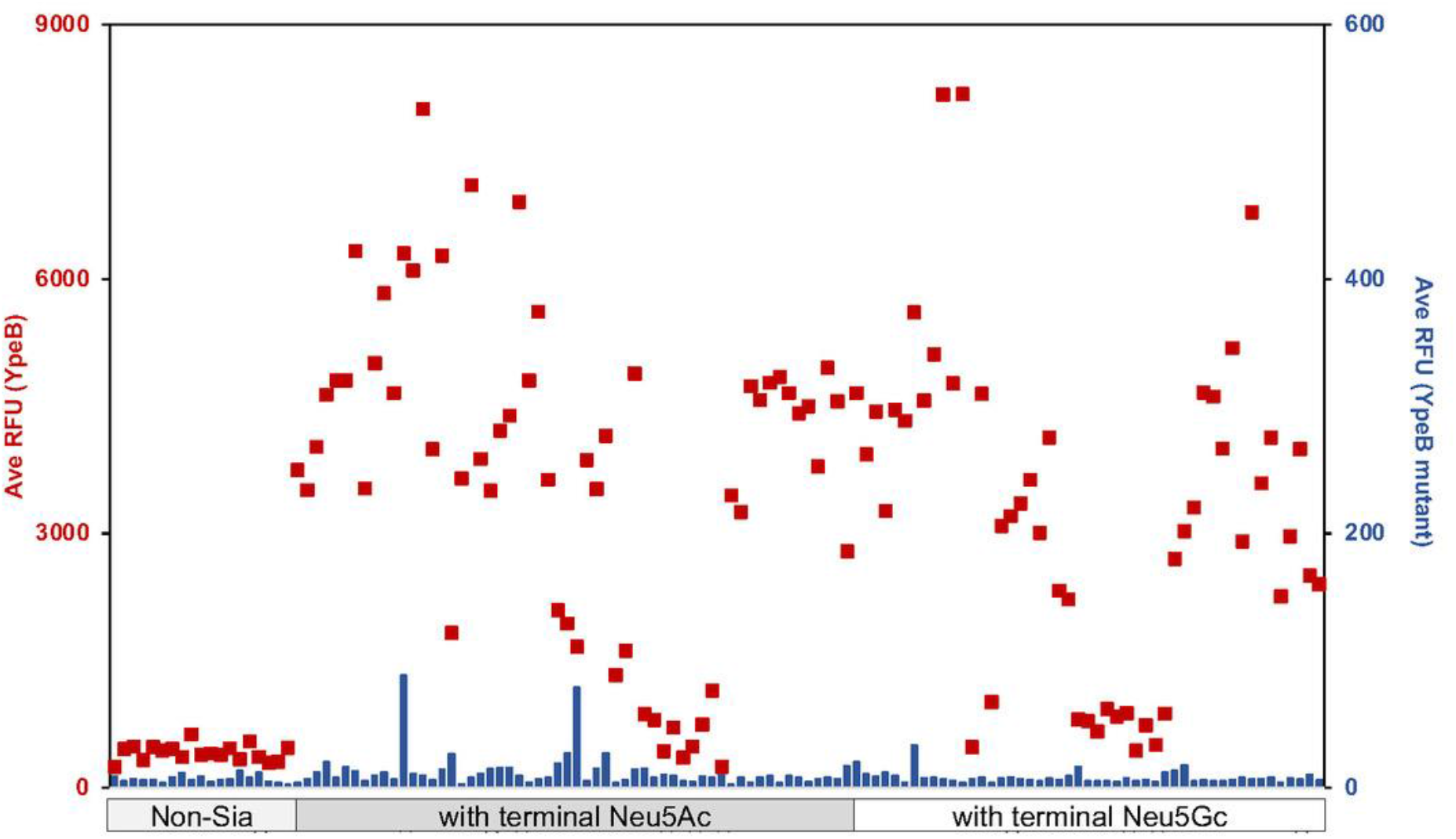
Site directed mutagenesis (Y77F) of YpeB eliminates sialoglycan binding. The red square dots showed Average RFU of YpeB binding to sialoglycans and blue bars showed average RFU of mutant YpeB binding to sialoglycans.

**Fig. 6.**
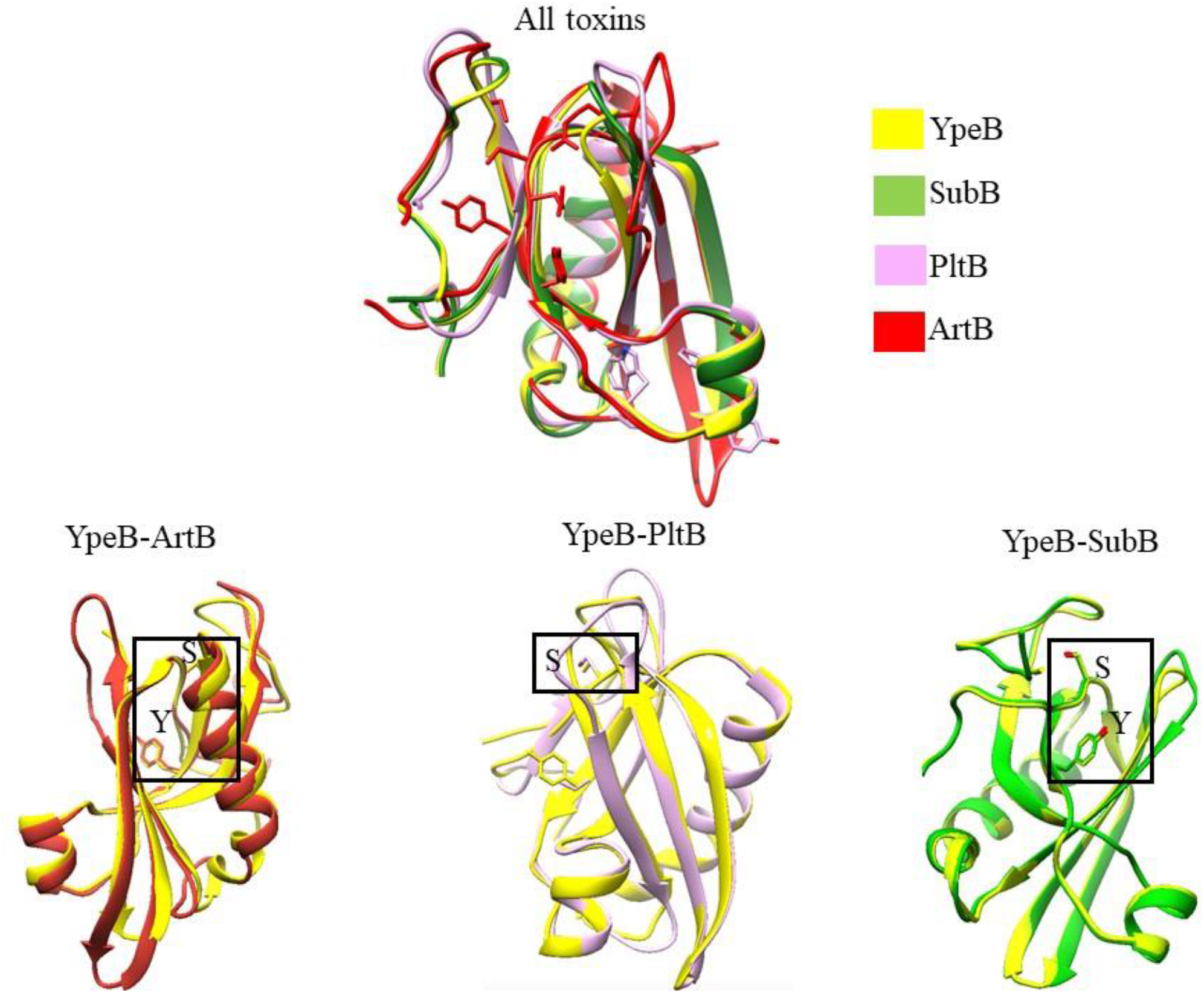
Comparative predicted structural modelling of three toxins (PltB, SubB, ArtB) with respect to YpeB. The box represents the location of critical Serine (S) and Tyrosine (Y) in each toxin comparative model. Both Tyrosine and Serine are present in three out of four toxin B subunits. Critical Tyrosine for Neu5Gc recognition is not present in *S*. Typhi PltB toxin.

### YpeB-induces apoptosis or necrosis in different cell lines, PBMCs and Splenocytes

In typical AB_5_ bacterial toxins, the B subunits mediate glycan binding, and the cognate A subunit then mediates toxicity. If so, the question arises as to why *Y. pestis* has an evolutionarily conserved isolated B subunit (A subunit is lacking), and what the role of this orphan B subunit might be. To ask whether the YpeB subunit without its counterpart can cause toxicity, general cytotoxicity was assessed by determining rates of necrosis in U937 human leukemia cells and COS-7 cells (from green monkey), exposed to 0.5 or 1.0 μg/ml YpeB for 16 hrs (Fig. 7A). Human PBMC and Splenocytes from WT mice were also incubated with different concentrations of YpeB (1-20 μg/ml) for 4 hrs (Fig. 7B). Since PBMC and mouse splenocytes are primary cell lines, we decreased the incubation time as well as using higher YpeB concentrations than for the cell lines. U-937 and COS-7 cell lines showed increased toxicity with increasing dose of YpeB. Similarly, PBMC and splenocytes showed increased toxicity with increased YpeB concentration. It is likely that the previously reported toxic/inflammatory effects of PltB of typhoid toxin (referred to as ArtB) on various cell types (26, 31) have an analogous mechanistic basis.

**Fig. 7.**
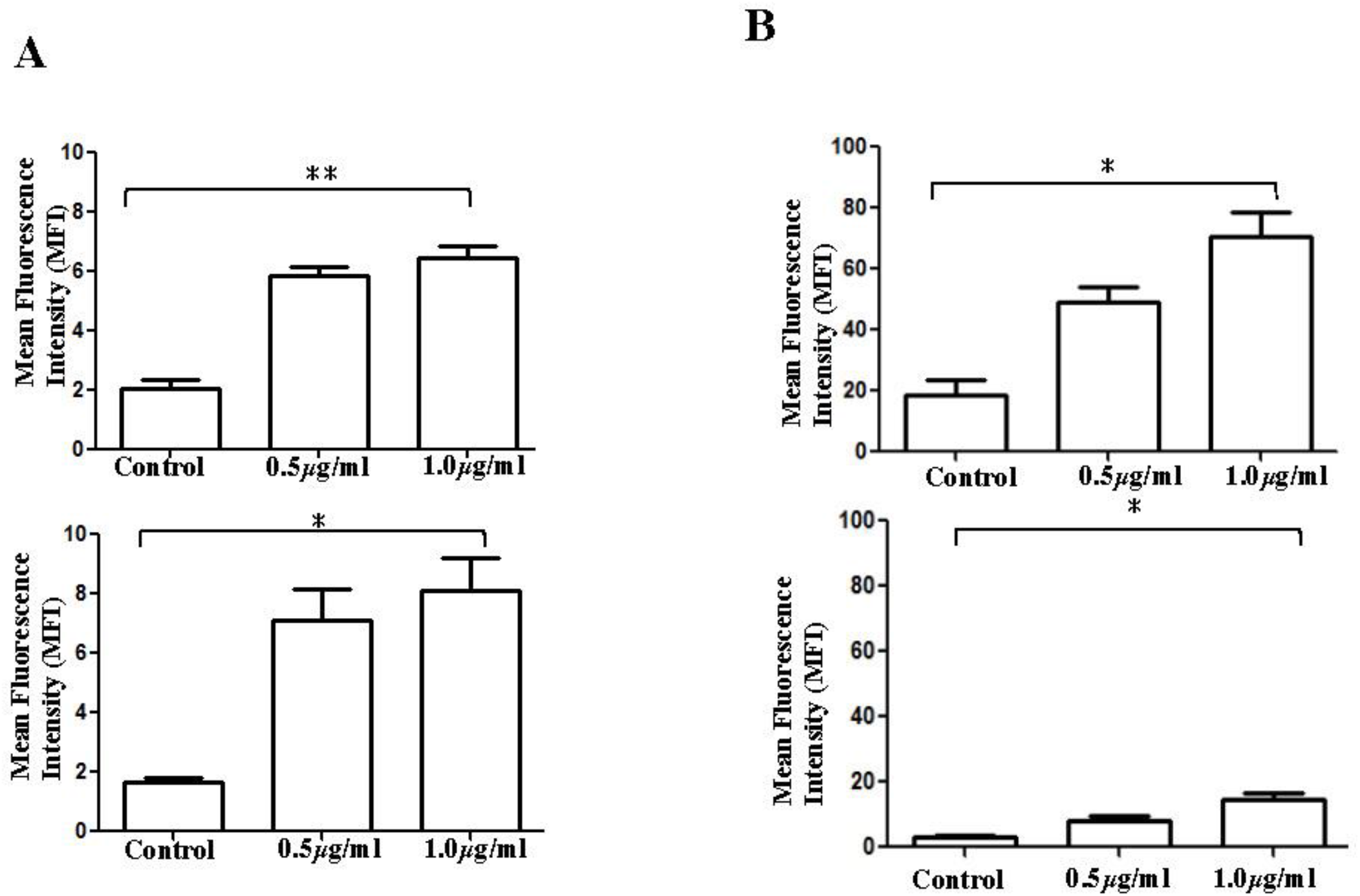
A. Effect of YpeB on cell lines. COS-7 cells (1×10^6^ cells) were incubated in DMEM medium with YpeB (0.5 μg/ml and 1.0μg/ml) for 16hr, then labelled with propidium iodide (PI) and analyzed by flow cytometry (Upper Panel). U-937 cells (1×10^6^ cells) were incubated in RPMI medium with YpeB (0.5 μg/ml and 1.0μg/ml) for 16hr, then labelled with propidium iodide (PI) and analyzed by flow cytometry (Lower panel). The data are represented as MFI (Mean Fluorescence Intensity) for each cell lines. ** P<0.01, *P<0.05. B. Effect of YpeB on fresh mouse Splenocytes and human peripheral blood mononuclear cells (PBMCs). Cells were incubated with YpeB (0.5 μg/ml and 1.0μg/ml) for 4hr and labelled with propidium iodide (PI) and analyzed by flow cytometry. The data are represented as MFI (Mean Fluorescence Intensity) for Splenocytes and PBMC. ** P<0.01, *P<0.05. (Since primary cells are more sensitive, we have reduced incubation time to 4hr).

### Mechanism of YpeB toxicity likely involves lectin-like crosslinking of glycoconjugates

CHOK1 cells were treated with active or inactive YpeB for 4 hrs, followed by measurement of cell viability using a MTT assay over the next 72 hrs. At low concentrations of YpeB (0.6 μg/ml, 1.25 μg/ml), the toxin activates cells to proliferate, whereas, at higher concentrations (2.5 μg/ml, 5.0 μg/ml), the cells were killed (Fig. 8A), likely reflecting a classic lectin-like crosslinking of glycoconjugates. After reaching saturation, the cells died because they were grown in very low FBS in the culture media. The activation at low concentration and killing at high concentration were not observed with inactive YpeB (Fig. 8B).

**Fig. 8.**
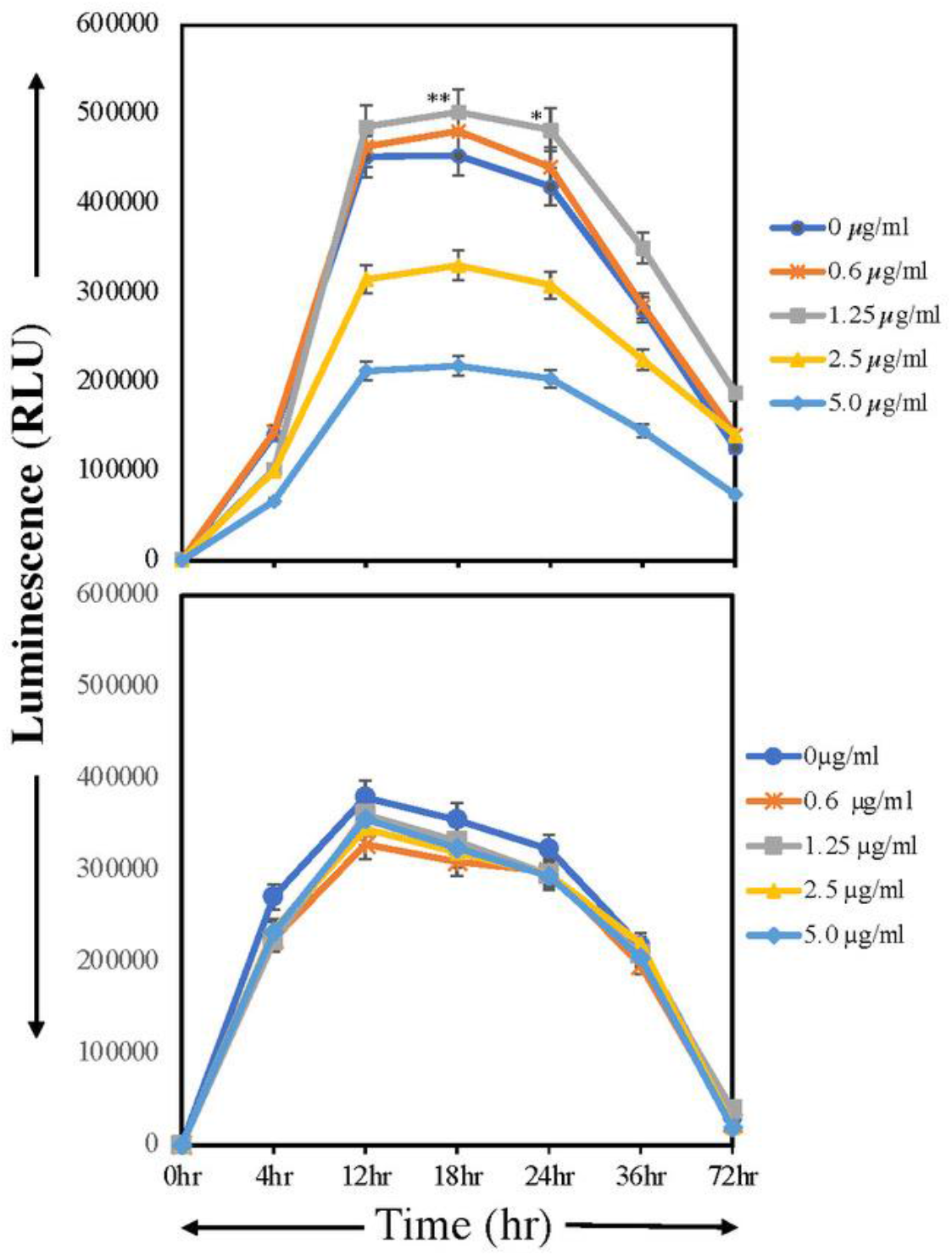
Cell growth and viability of CHOK1 cells (5000 cells/well) with active (Upper Panel) or non-binding (Lower panel) YpeB toxin B subunit. Cells were incubated with toxin concentrations ranging from 0.6-5.0 μg/ml at different time points ranging from 0-72hr using MTT cell viability assay on ELISA plate reader at 570 nm (described in experimental procedures). The upper panel shows lectin like stimulation, whereas no stimulation and proliferation were observed in lower panel, where cells incubated with mutant YpeB (serum free condition and toxin added after that). ** P<0.01, *P<0.05.

### Subcutaneous injection of YpeB toxin causes proliferation of B cells

Several studies have reported inflammatory responses initiated by the host immune system in response to Yersinia pestis. Among them is the hyperproliferation of B cells (32). When YpeB was injected subcutaneously, examination of spleen sections showed proliferation of B cells marked with B220 (B cell marker) (Fig. 9). This is alsolikely due to the lectin-like properties of YpeB.

**Fig. 9.**
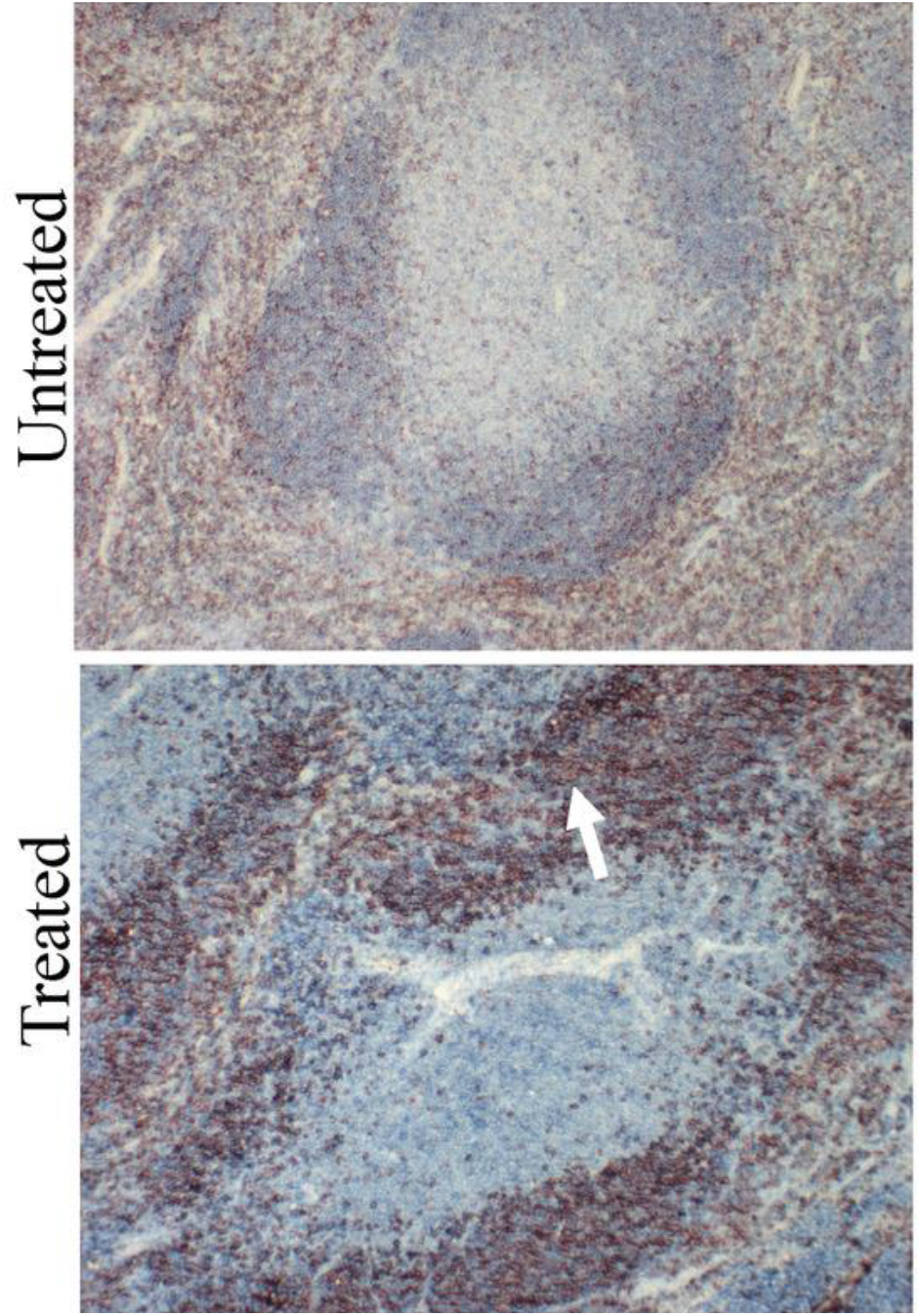
Subcutaneous injection of YpeB causes proliferation of B cells in spleen sections. The upper panel is the spleen section untreated with YpeB and lower panel shows YpeB treated spleen section. Dark brown dots in YpeB-treated spleen section shows proliferation of B cells (stained with B220)

## Conclusions and Perspectives

Host specificity of pathogens is the result of various factors including genetic diversity, physiological conditions, and ecological opportunities. There are two general categories of bacterial pathogens: one group known as specialists which establish intimate relationships with single (or a closely related group of) hosts, whereas the other group known as generalists that are capable of infecting a wide range of hosts. Most pathogens exhibit some degree of host preference, for example in malaria *Plasmodium falciparum* is specific for humans, whereas *Plasmodium reichenowi* specifically infects chimpanzees (33, 34).

The features that govern host specificity are not yet fully understood. In the present study, we have explored the evolutionary relationship between A and B subunits of bacterial AB_5_ toxins. Due to the lack of evolutionary relationship and independent evolution of these two subunits it is evident that B subunits are evolving based on cross species transmission between bacteria that infect a wide range of hosts.

In the present study we have focused attention on *Y. pestis* YpeB, which is clearly a member of the AB_5_ toxin B subunit family. Among eleven species of Yersinia, only three are pathogenic. Among them the deadliest is *Y. pestis*, the causative agent of plague, which was responsible for the black death of Europe in the 14th century (35). YpeB shares strongest sequence similarity and phylogenetic relationships with *E. coli* SubB and *S*. Typhi PltB. Unlike SubB and PltB, YpeB binds to both Neu5Ac and Neu5Gc terminated glycans as previously discovered for *S*. Typhimurium ArtB (7). The broad glycan binding pattern of YpeB is consistent with a broad range of host specificity of a pathogen that can infect a wide variety of animals ranging from mouse, rats, rabbits, cats, dogs, squirrels to humans (36). In terms of severity of diseases, *S*. Typhimurium causes mild gastrointestinal infections in humans (37), whereas *Y. pestis* has been involved in three human plague pandemics (38).

A comparison of *YpeB* and other toxin B subunits (*PltB, SubB and ArtB*) glycan binding patterns reveals differences in binding to host glycans reflecting the host range, virulence and pathogenesis of the bacteria that produce them. PltB exhibits exquisite preference for human-specific Neu5Ac-terminated sialoglycans on surface glycoproteins that serve as its receptors (4). In contrast, SubB, prefers to bind to Neu5Gc-terminated sialoglycans on surface glycoproteins that serve as its receptors (5). It is known that purified SubAB is highly lethal for mice when injected intraperitoneally, causing pathology similar to that seen in human cases of hemolytic uremic syndrome (14). This is consistent with preferential expression of Neu5Gc by rodents. Strains of *E. coli* producing SubAB have also been associated with severe disease in humans, including hemolytic uremic syndrome (10, 28). However, these strains also produce Shiga toxin (StxAB), so the contribution of SubAB to human disease/pathology remains uncertain. The binding preference of ArtB for both Neu5Ac and Neu5Gc terminated glycans is entirely consistent with the broad host range of the bacterium that harbors it. Interestingly, despite having broad glycan binding patterns, YpeB lacks binding with 4-O-acetylated Neu5Ac and Neu5Gc glycans. Considering high expression of 4-O-acetylated glycans in horses (39), it is interesting that they are anecdotally noted to be less affected in plague epidemics. In the following paper (Sasmal et al. 2020), we describe another orphan B subunit toxin from Yersinia species (*Yersinia enterocolitica* (YenB)) with an even broader range of Sia recognition than YpeB of *Yersinia pestis*, again correlating with a broader range of hosts.

Unlike other AB_5_ toxin members, another interesting feature about YpeB is that, without an enzymatically active A subunit, the B subunit alone causes toxicity to different cell types. The cytotoxic effect of YpeB is likely based on binding to the target cells and a lectin-like action of the protein (40), as the Y77F glycan binding site of mutant YpeB has no toxic effects. Toxicity was highest in mouse splenocytes as compared to human cells (cells lines and PBMC) and monkey kidney cell lines. The higher toxicity in mouse splenocytes could be due to higher concentration of Neu5Ac- and Neu5Gc-terminating glycans (41) as compared to green monkey kidney cells. In contrast, human cells showed toxicity owing to their dominant Neu5Ac.

With the help of the already known structure of SubB, structural modellling of YpeB reveals conserved amino acids essential for host glycan binding. For *E. coli* SubB, it is evident that Ser12 and Tyr78 are essential for binding to sialoglycans. In SubB Tyr78 which interacts with the extra OH group of Neu5Gc and is thus critical for binding to Neu5Gc, but does not impact the albeit weaker recognition of Neu5Ac (5). Interestingly, the analogous Y77F mutation in YpeB abolished the binding to both Neu5Ac and Neu5Gc glycans, which indicates that the binding tyrosine in YpeB plays a more prominent role in glycan binding than it does in SubB.

Since we do not know how the B subunit alone causes toxicity without an enzymatically active A subunit, we speculate that multivalent YpeB behaves like a lectin that activates the target cell by crosslinking cell surface glycans, thereby exerting multifaceted effects such as signal modulation, membrane reshuffling, cell-cell adhesions and thus contributes to the pathogenic property of the bacterium (42). Significantly, the non-sialic acid binding mutant of YpeB Y77F showed reduced toxicity and cell. We have shown that YpeB *exerts* toxicity only at relatively high concentrations. A similar case was observed for wheat germ agglutinin (WGA), high concentrations of which kills cells (43). In addition, higher doses of *S*. Typhi PltB causes inflammation and vacuolation with less incubation time (31).

Taken together, our study is the first to report the broad range of glycan binding specificity of YpeB that binds to both Neu5Ac and Neu5Gc terminated glycans (except 4-OAc glycans), which is concordant with the broad host range of *Y. pestis*. The orphan B subunit causes toxicity in different cell types of human, mouse and monkey origin, and behaves like a lectin to induce toxicity in cells. Nevertheless, more work is required to understand why YpeB did not bind to 4-OAc glycans, which are highly expressed in horses, that are speculated to be resistant to plague.

## Experimental Procedures

### Phylogenetic inference of AB_5_ toxins among species

The genome sequence of 33 bacterial species were retrieved via BLAST from NCBI (www.ncbi.nlm.nih) and Uniprot, using as queries A and B subunits of the AB_5_ family toxins from *S*. Typhi (PltAB) and *E. coli* (SubAB Sequence alignments were performed using Clustal W implemented in MEGAv7.0 (44). Based on sequence retrieved from 33 species, a neighbor-joining phylogenetic tree was constructed. All ambiguous positions were removed for each sequence pair (pairwise deletion option). The sequence of A subunits was retrieved following the above method. Since it is known that some bacterial species lacks active A subunit and the phylogenetic tree based on A subunit comprised of bacterial species that contains A subunit. Additionally, sequences of 16S rRNA from each species were retrieved and neighbor-joining (NJ) phylogenetic tree was constructed using MEGAv.0 7(44). Accession number of the retrieved sequences were added on the phylogenetic tree.

### Purification of YpeB

The *Y. pestis ypeB* open reading frame (ORF) was chemically synthesized (GenScript) incorporating flanking 5’ and 3’ *Eco*RI and *Hin*dIII restriction sites, as well as a region encoding a His_6_ tag on the 3’ terminus of *YpeB*, and was cloned into pBAD18, so that the ORF is under the control of the vector *ara* promoter. The construct was then transformed into an *E. coli* BL21(DE3) *lpxM* mutant that produces a penta-acylated lipopolysaccharide (LPS) that has very low endotoxic activity. The recombinant bacterium was grown in 500 ml LB Broth at 37°C to late logarithmic phase, diluted 50:50 with fresh medium supplemented with 0.2% (w/v) arabinose to induce *YpeB* expression, and then incubated overnight at 26°C. Cells were harvested by centrifugation and re-suspended in 20 ml loading buffer (50 mM sodium phosphate, 300 mM NaCl, pH 8.0), and lysed with a sonicator for 10 min (On/Off). Cell debris was removed by centrifugation at 40,000 × *g* for 30 min at 4°C. The supernatant was then loaded onto a preequilibrated 2-ml Ni-NTA column. The Ni-NTA column was washed with 40 ml loading buffer (mentioned above), and bound proteins were eluted with a 30-ml gradient of 0 to 500 mM imidazole in loading buffer. Ten tubes of 3-ml fractions were collected and analyzed by SDS-PAGE, followed by staining with Coomassie blue or Western blotting with monoclonal anti-His_6_ (Genscript, USA). YpeB migrated as a single ~16-KDa band on SDS-PAGE when stained with Coomassie.

### Sialoglycan Microarray

The sialoglycan microarray method was adapted and modified from the literature reported earlier (45, 46). Defined sialosides with amine linker were chemoenzymatically synthesized and then quantitated utilizing an improved DMB-HPLC method. 100 μM of sialoglycan solution (in 300mM Na-phosphate buffer, pH 8.4) was printed in quadruplets on NHS-functionalized glass slides (PolyAn 3D-NHS; catalog# PO-10400401) using an ArrayIt SpotBot^®^ Extreme instrument. The slides were blocked (0.05M ethanolamine solution in 0.1M Tris-HCl, pH 9.0), washed with warm Milli-Q water and dried. The slides were then fitted in a multi-well microarray hybridization cassette (ArrayIt, CA) to divide into 8 wells and rehydrated with 400 μl of Ovalbumin (1% w/v, PBS) for one hour in a humid chamber with gentle shaking. After that the blocking solution was discarded and a 400 μl of solution of the toxin B-subunit (30 μg/ml) in the same blocking buffer was added to the individual well. The slides were incubated for 2h at room temperature with gentle shaking and the slides were washed with PBS-Tween (0.1% v/v). The wells were then treated with Cy3-conjugated anti-His (Rockland Antibodies & Assays; Cat# 200-304-382) secondary antibody at 1:500 dilution in PBS. Followed by gentle shaking for 1 hour in the dark and humid chamber. The slides were then washed, dried, and scanned with a Genepix 4000B scanner (Molecular Devices Corp., Union City, CA) at wavelength 532 nm. Data analysis was performed using the Genepix Pro 7.3 software (Molecular Devices Corp., Union City, CA). The raw data analysis and sorting using the numerical codes were performed on Microsoft Excel. Local background subtraction was performed, and data were plotted separately for each subarray. The binding specificity to glycoconjugates for each protein was plotted based on the average RFU (Relative Fluorescence Units) versus glycan IDs.

### Cell Culture and Monosaccharide Supplementation

In order to determine the sialic acid dependent binding of YpeB toxin B subunit we used the cultured cell line LEC29.Lec32. These CHO cells (that lack sialic acid production due to mutation in CMAS gene (30). LEC29.Lec32 cells was propagated as an adherent cell lines supplemented with alpha MEM medium with 10% (v/v) heat-inactivated fetal calf serum, 2 mM glutamine, 100 U of penicillin per mL, and 100 μg of streptomycin per mL in a humidified 5% CO_2_, 37 °C atmosphere. For medium supplementation, Neu5Ac (3mM) and Neu5Gc (3mM) were dissolved in PBS, titrated to a neutral pH, and filter sterilized. The sugars were added at the indicated concentrations. For the sugar turnover experiments, the cells were fed as mentioned above and on day 0, the cell culture medium was switched to respective medium, supplemented with 1% (v/v) Nutridoma (Roche), 100 U of penicillin per mL, and 100 μg of streptomycin without Neu5Ac and Neu5Gc.

### Flow Cytometry Analysis

Cells were washed with PBS and incubated with different concentration of YpeB in PBS and incubated with Alexa Fluor 647 anti-His antibody for 30 min on ice. After an additional washing step, the cells were analyzed by Flow Cytometry.

### Cell culture and toxin treatment

To understand the toxicity of B subunit in cells, we cultured different cell lines: Cos-7 cells (African Green monkey kidney fibroblast), grown in DMEM supplemented with 10% FCS and 50 IU of penicillin/50 μg/ml of streptomycin and U937 (human monocytes) cells, grown in RPMI 1640 medium supplemented with 10 mM HEPES, 2 mM L-glutamine, 1 mM sodium pyruvate, 10% (v/v) heat-inactivated FCS, 50 IU of penicillin/ 50 mg/ml streptomycin, at 37°C and 5% CO_2_. For the toxin treatment, cells were seeded into 24-well tissue culture plates; (1 × 10^6^/well) and were exposed to YpeB toxin B subunit at the indicated concentration in 300 μl of culture medium. Additionally, PBMC and mouse spleenocytes were also cultured in RPMI 1640 medium supplemented with 10 mM HEPES, 2 mM L-glutamine, 1 mM sodium pyruvate, 10% (v/v) heat-inactivated FCS, 50 IU of penicillin/ 50 mg/ml streptomycin and treated with indicated concentration of toxin at different time interval and visualized under FACSCalibur.

### PBMC isolation and Splenocytes from WT mice

Freshly drawn venous blood was used to isolate PBMC with Ficoll hypaque method. After isolating PBMCs, cells were washed with PBS and centrifuged for 10 min at 200 x *g*. After washing, the supernatant was discarded, and the cells were treated with ACK lysis buffer and incubate for 10 min. This step repeated till the trace of red blood cells vanished. The cells were counted with hemocytometer and approximately ~2×10^6^ cells were seeded in RPMI+10% (v/v) FCS in a tissue culture plate and incubated with toxins at different concentration. The similar steps were followed for Splenocytes the freshly isolated spleens were crushed using plunger end of the syringe and filtered through the cell strainer into the petri dish. The single cell suspension was centrifuged and washed three times for ten min at 200g. The cells were treated with ACK lysis buffer and incubated for ten min. After ACK lysis, the cells were suspended in RPMI+10% FCS and seeded in a cell culture plate and incubated with 1ug/ml and 10ug/ml toxin concentration.

### Examination of apoptosis and necrosis

Exploring the role of B subunit in toxicity, cultured cells were stained with markers of apoptosis as well as necrosis. Cells were detected by differential staining with annexin V (which stains apoptotic and necrotic cells) and propidium iodide (PI) (which stains necrotic cells only) using a Annexin-V Apoptosis Kit 2 (Invitrogen V13241) according to the manufacturer’s instructions. Briefly, 10^6^ cells were treated for 16 hr with 10 μg/ml YpeB and transferred from the wells of the tissue culture plate into fluorescence-activated cell sorter (FACS) tubes (Falcon 352008). The cells were washed three times with 3 ml annexin V binding buffer and resuspended in 50 μl of annexin V working reagent, consisting of 1/25 annexin V-Alexa 488 (which emits at 519 nm) and 1 μg/ml PI (which emits at 617 nm) in annexin V binding buffer. The cells were incubated at room temperature in the dark for 30 min and then washed and analyzed immediately on a FACSCalibur flow cytometer.

### Structural modelling

Since YpeB toxin B subunit is a novel toxin without a counterpart A-subunit, the crystal structure through X-ray crystallography is a challenge. The structure of the YpeB was predicted using PyMol software. The highest scored structure was selected for the structural comparisons, in this case we selected Ecoli SubB subunit (PDB ID: 3DWA), because of its high similarity. The sialic acid binding site was predicted after docking it with Neu5Gc. In addition, comparison of structures of ArtB, PltB and SubB with YpeB were done using Chimera software (47).

TheYpeB amino acid sequenceswere aligned using the program EXPRESSO (48) and homology modeling was performed employing MODELLER software (49), using the crystal structure of SubB (3DWP) as a template.

### Cell Viability Assay

Cell viability assays were performed in CHOK1 cell lines. Promega MTT, absorbed into the cell and eventually the mitochondria, is broken down into formazan by mitochondria succinate dehydrogenase. Accumulation of formazan reflects the activity of mitochondria directly and cell viability indirectly. Cell viability was measured by the MTT assay. CHOK1 cells were seeded on 96-well plates at a density of 5×10^3^ cells/well, cultured overnight in advanced media with 2% FBS. Different concentrations of YpeB active and inactive toxin B subunits were added to the wells and incubated for 4 hrs. In addition, the plant lectin MAL-1 was also added because it lacks the active component (like YpeB) and acts as a control. After incubation, MTT was added at a concentration of 0.5 mg/ml after medium (200 μl) was added to each well. The plates were incubated at 37°C for different time points (0,4hr,12hr,24hr,72hr) and was measured on an ELISA reader (Bio-Rad, Hercules, CA, USA) at a wavelength of 570 nm and O.D was recorded.

### Histopathology

To understand the role of YpeB in toxicity and B cell proliferation, in vivo, we injected YpeB subcutaneously into flanks of WT mice, which were euthanized and sacrificed after 3 days. Spleens were harvested and frozen for frozen sections and immuno-stained using several markers including the proliferative marker (B220) for B cells.

### Statistical analysis

All data was analyzed with GraphPad Prism version 6.0 (GraphPad Software, San Diego, CA) and comparisons were made by Student’s *t* test (two-tailed), one-way analysis of variance or two-way analysis of variance, as appropriate. A probability of *P* < 0.05 indicated as a statistical significance.

## Acknowledgments

This work was supported by NIH Grant R01GM32373 (to A.V.).

## Conflicts of Interest

The authors declare no competing interests.

## REFERENCES

1. Beddoe, T., Paton, A. W., Le Nours, J., Rossjohn, J. and Paton, J. C. (2010) Structure, biological functions and applications of the AB5 toxins. Trends Biochem Sci 35, 411–418

2. Ng, N., Littler, D., Le Nours, J., Paton, A. W., Paton, J. C., Rossjohn, J. and Beddoe, T. (2013) Cloning, expression, purification and preliminary X-ray diffraction studies of a novel AB5 toxin. Acta Crystallogr Sect F Struct Biol Cryst Commun 69, 912–915

3. Paton, A. W., Beddoe, T., Thorpe, C. M., Whisstock, J. C., Wilce, M. C., Rossjohn, J., Talbot, U. M. and Paton, J. C. (2006) AB5 subtilase cytotoxin inactivates the endoplasmic reticulum chaperone BiP. Nature 443, 548–552

4. Deng, L., Song, J., Gao, X., Wang, J., Yu, H., Chen, X., Varki, N., Naito-Matsui, Y., Galán, J. E. and Varki, A. (2014) Host adaptation of a bacterial toxin from the human pathogen Salmonella Typhi. Cell 159, 1290–1299

5. Byres, E., Paton, A. W., Paton, J. C., Löfling, J. C., Smith, D. F., Wilce, M. C., Talbot, U. M., Chong, D. C., Yu, H., Huang, S., Chen, X., Varki, N. M., Varki, A., Rossjohn, J. and Beddoe, T. (2008) Incorporation of a non-human glycan mediates human susceptibility to a bacterial toxin. Nature 456, 648–652

6. Song, J., Gao, X. and Galán, J. E. (2013) Structure and function of the Salmonella Typhi chimaeric A(2)B(5) typhoid toxin. Nature 499, 350–354

7. Gao, X., Deng, L., Stack, G., Yu, H., Chen, X., Naito-Matsui, Y., Varki, A. and Galán, J. E. (2017) Evolution of host adaptation in the Salmonella typhoid toxin. Nat Microbiol 2, 1592–1599

8. van den Akker, F., Sarfaty, S., Twiddy, E. M., Connell, T. D., Holmes, R. K. and Hol, W. G. (1996) Crystal structure of a new heat-labile enterotoxin, LT-IIb. Structure 4, 665–678

9. Locht, C. and Antoine, R. (1995) A proposed mechanism of ADP-ribosylation catalyzed by the pertussis toxin S1 subunit. Biochimie 77, 333–340

10. Paton, A. W., Srimanote, P., Talbot, U. M., Wang, H. and Paton, J. C. (2004) A new family of potent AB(5) cytotoxins produced by Shiga toxigenic Escherichia coli. J Exp Med 200, 35–46

11. Ng, N. M., Littler, D. R., Paton, A. W., Le Nours, J., Rossjohn, J., Paton, J. C. and Beddoe, T. (2013) EcxAB is a founding member of a new family of metalloprotease AB5 toxins with a hybrid cholera-like B subunit. Structure 21, 2003–2013

12. Naha, A., Mandal, R. S., Samanta, P., Saha, R. N., Shaw, S., Ghosh, A., Chatterjee, N. S., Dutta, P., Okamoto, K., Dutta, S. and Mukhopadhyay, A. K. (2020) Deciphering the possible role of ctxB7 allele on higher production of cholera toxin by Haitian variant Vibrio cholerae O1. PLoS Negl Trop Dis 14, e0008128

13. Sack, D. A., Sack, R. B., Nair, G. B. and Siddique, A. K. (2004) Cholera. Lancet 363, 223–233

14. Wang, H., Paton, J. C. and Paton, A. W. (2007) Pathologic changes in mice induced by subtilase cytotoxin, a potent new Escherichia coli AB5 toxin that targets the endoplasmic reticulum. J Infect Dis 196, 1093–1101

15. Chou, H. H., Takematsu, H., Diaz, S., Iber, J., Nickerson, E., Wright, K. L., Muchmore, E. A., Nelson, D. L., Warren, S. T. and Varki, A. (1998) A mutation in human CMP-sialic acid hydroxylase occurred after the Homo-Pan divergence. Proc Natl Acad Sci U S A 95, 11751–11756

16. Perry, R. D. and Fetherston, J. D. (1997) Yersinia pestis--etiologic agent of plague. Clin Microbiol Rev 10, 35–66

17. Steensma, D. P. and Kyle, R. A. (2020) Alexandre Yersin: Discoverer of the Plague Bacillus. Mayo Clin Proc 95, e7–e8

18. Dziarski, R. (2006) Deadly plague versus mild-mannered TLR4. Nat Immunol 7, 1017–1019

19. Bevins, S. N., Baroch, J. A., Nolte, D. L., Zhang, M. and He, H. (2012) Yersinia pestis: examining wildlife plague surveillance in China and the USA. Integr Zool 7, 99–109

20. Anisimov, A. P. and Amoako, K. K. (2006) Treatment of plague: promising alternatives to antibiotics. J Med Microbiol 55, 1461–1475

21. Morelli, G., Song, Y., Mazzoni, C. J., Eppinger, M., Roumagnac, P., Wagner, D. M., Feldkamp, M., Kusecek, B., Vogler, A. J., Li, Y., Cui, Y., Thomson, N. R., Jombart, T., Leblois, R., Lichtner, P., Rahalison, L., Petersen, J. M., Balloux, F., Keim, P., Wirth, T., Ravel, J., Yang, R., Carniel, E. and Achtman, M. (2010) Yersinia pestis genome sequencing identifies patterns of global phylogenetic diversity. Nat Genet 42, 1140–1143

22. Rasmussen, S., Allentoft, M. E., Nielsen, K., Orlando, L., Sikora, M., Sjögren, K. G., Pedersen, A. G., Schubert, M., Van Dam, A., Kapel, C. M., Nielsen, H. B., Brunak, S., Avetisyan, P., Epimakhov, A., Khalyapin, M. V., Gnuni, A., Kriiska, A., Lasak, I., Metspalu, M., Moiseyev, V., Gromov, A., Pokutta, D., Saag, L., Varul, L., Yepiskoposyan, L., Sicheritz-Pontén, T., Foley, R. A., Lahr, M. M., Nielsen, R., Kristiansen, K. and Willerslev, E. (2015) Early divergent strains of Yersinia pestis in Eurasia 5,000 years ago. Cell 163, 571–582

23. Cui, Y., Yu, C., Yan, Y., Li, D., Li, Y., Jombart, T., Weinert, L. A., Wang, Z., Guo, Z., Xu, L., Zhang, Y., Zheng, H., Qin, N., Xiao, X., Wu, M., Wang, X., Zhou, D., Qi, Z., Du, Z., Wu, H., Yang, X., Cao, H., Wang, H., Wang, J., Yao, S., Rakin, A., Li, Y., Falush, D., Balloux, F., Achtman, M., Song, Y., Wang, J. and Yang, R. (2013) Historical variations in mutation rate in an epidemic pathogen, Yersinia pestis. Proc Natl Acad Sci U S A 110, 577–582

24. Gage, K. L., Dennis, D. T., Orloski, K. A., Ettestad, P., Brown, T. L., Reynolds, P. J., Pape, W. J., Fritz, C. L., Carter, L. G. and Stein, J. D. (2000) Cases of cat-associated human plague in the Western US, 1977-1998. Clin Infect Dis 30, 893–900

25. Yang, R. (2018) Plague: Recognition, Treatment, and Prevention. J Clin Microbiol 56,

26. Wang, H., Paton, J. C., Herdman, B. P., Rogers, T. J., Beddoe, T. and Paton, A. W. (2013) The B subunit of an AB5 toxin produced by Salmonella enterica serovar Typhi up-regulates chemokines, cytokines, and adhesion molecules in human macrophage, colonic epithelial, and brain microvascular endothelial cell lines. Infect Immun 81, 673–683

27. Padler-Karavani, V., Song, X., Yu, H., Hurtado-Ziola, N., Huang, S., Muthana, S., Chokhawala, H. A., Cheng, J., Verhagen, A., Langereis, M. A., Kleene, R., Schachner, M., de Groot, R. J., Lasanajak, Y., Matsuda, H., Schwab, R., Chen, X., Smith, D. F., Cummings, R. D. and Varki, A. (2012) Cross-comparison of protein recognition of sialic acid diversity on two novel sialoglycan microarrays. J Biol Chem 287, 22593–22608

28. Paton, A. W., Woodrow, M. C., Doyle, R. M., Lanser, J. A. and Paton, J. C. (1999) Molecular characterization of a Shiga toxigenic Escherichia coli O113:H21 strain lacking eae responsible for a cluster of cases of hemolytic-uremic syndrome. J Clin Microbiol 37, 3357–3361

29. Saitoh, M., Tanaka, K., Nishimori, K., Makino, S. I., Kanno, T., Ishihara, R., Hatama, S., Kitano, R., Kishima, M., Sameshima, T., Akiba, M., Nakazawa, M., Yokomizo, Y. and Uchida, I. (2005) The artAB genes encode a putative ADP-ribosyltransferase toxin homologue associated with Salmonella enterica serovar Typhimurium DT104. Microbiology (Reading) 151, 3089–3096

30. Potvin, B., Raju, T. S. and Stanley, P. (1995) Lec32 is a new mutation in Chinese hamster ovary cells that essentially abrogates CMP-N-acetylneuraminic acid synthetase activity. J Biol Chem 270, 30415–30421

31. Herdman, B. P., Paton, J. C., Wang, H., Beddoe, T. and Paton, A. W. (2017) Vacuolation Activity and Intracellular Trafficking of ArtB, the Binding Subunit of an AB5 Toxin Produced by Salmonella enterica Serovar Typhi. Infect Immun 85,

32. Hinnebusch, B. J., Jarrett, C. O., Callison, J. A., Gardner, D., Buchanan, S. K. and Plano, G. V. (2011) Role of the Yersinia pestis Ail protein in preventing a protective polymorphonuclear leukocyte response during bubonic plague. Infect Immun 79, 4984–4989

33. Martin, M. J., Rayner, J. C., Gagneux, P., Barnwell, J. W. and Varki, A. (2005) Evolution of human-chimpanzee differences in malaria susceptibility: relationship to human genetic loss of N-glycolylneuraminic acid. Proc Natl Acad Sci U S A 102, 12819–12824

34. Varki, A. and Gagneux, P. (2009) Human-specific evolution of sialic acid targets: explaining the malignant malaria mystery. Proc Natl Acad Sci U S A 106, 14739–14740

35. Heroven, A. K. and Dersch, P. (2014) Coregulation of host-adapted metabolism and virulence by pathogenic yersiniae. Front Cell Infect Microbiol 4, 146

36. Oyston, P. C. and Williamson, D. (2011) Plague: Infections of Companion Animals and Opportunities for Intervention. Animals (Basel) 1, 242–255

37. Chaudhuri, D., Roy Chowdhury, A., Biswas, B. and Chakravortty, D. (2018) Salmonella Typhimurium Infection Leads to Colonization of the Mouse Brain and Is Not Completely Cured With Antibiotics. Front Microbiol 9, 1632

38. Wagner, D. M., Klunk, J., Harbeck, M., Devault, A., Waglechner, N., Sahl, J. W., Enk, J., Birdsell, D. N., Kuch, M., Lumibao, C., Poinar, D., Pearson, T., Fourment, M., Golding, B., Riehm, J. M., Earn, D. J., Dewitte, S., Rouillard, J. M., Grupe, G., Wiechmann, I., Bliska, J. B., Keim, P. S., Scholz, H. C., Holmes, E. C. and Poinar, H. (2014) Yersinia pestis and the plague of Justinian 541-543 AD: a genomic analysis. Lancet Infect Dis 14, 319–326

39. Wasik, B. R., Barnard, K. N., Ossiboff, R. J., Khedri, Z., Feng, K. H., Yu, H., Chen, X., Perez, D. R., Varki, A. and Parrish, C. R. (2017) Distribution of O-Acetylated Sialic Acids among Target Host Tissues for Influenza Virus. mSphere 2,

40. Mendelsohn, J., Skinner, A. and Kornfeld, S. (1971) The rapid induction by phytohemagglutinin of increased alpha-aminoisobutyric acid uptake by lymphocytes. J Clin Invest 50, 818–826

41. Ji, S., Wang, F., Chen, Y., Yang, C., Zhang, P., Zhang, X., Troy, F. A. and Wang, B. (2017) Developmental changes in the level of free and conjugated sialic acids, Neu5Ac, Neu5Gc and KDN in different organs of pig: a LC-MS/MS quantitative analyses. Glycoconj J 34, 21–30

42. Singh, R. S., Walia, A. K. and Kanwar, J. R. (2016) Protozoa lectins and their role in host-pathogen interactions. Biotechnol Adv 34, 1018–1029

43. Ryva, B., Zhang, K., Asthana, A., Wong, D., Vicioso, Y. and Parameswaran, R. (2019) Wheat Germ Agglutinin as a Potential Therapeutic Agent for Leukemia. Front Oncol 9, 100

44. Kumar, S., Stecher, G. and Tamura, K. (2016) MEGA7: Molecular Evolutionary Genetics Analysis Version 7.0 for Bigger Datasets. Mol Biol Evol 33, 1870–1874

45. Meng, C., Sasmal, A., Zhang, Y., Gao, T., Liu, C. C., Khan, N., Varki, A., Wang, F. and Cao, H. (2018) Chemoenzymatic Assembly of Mammalian O-Mannose Glycans. Angew Chem Int Ed Engl 57, 9003–9007

46. Lu, N., Ye, J., Cheng, J., Sasmal, A., Liu, C. C., Yao, W., Yan, J., Khan, N., Yi, W., Varki, A. and Cao, H. (2019) Redox-Controlled Site-Specific α2-6-Sialylation. J Am Chem Soc 141, 4547–4552

47. Pettersen, E. F., Goddard, T. D., Huang, C. C., Couch, G. S., Greenblatt, D. M., Meng, E. C. and Ferrin, T. E. (2004) UCSF Chimera--a visualization system for exploratory research and analysis. J Comput Chem 25, 1605–1612

48. O’Sullivan, O., Suhre, K., Abergel, C., Higgins, D. G. and Notredame, C. (2004) 3DCoffee: combining protein sequences and structures within multiple sequence alignments. J Mol Biol 340, 385–395

49. Eswar, N., Webb, B., Marti-Renom, M. A., Madhusudhan, M. S., Eramian, D., Shen, M. Y., Pieper, U. and Sali, A. (2006) Comparative protein structure modeling using Modeller. Curr Protoc Bioinformatics Chapter 5, Unit-5.6

50. Sporecke, I., Castro, D. and Mekalanos, J. J. (1984) Genetic mapping of Vibrio cholerae enterotoxin structural genes. J Bacteriol 157, 253–261

51. Fowler, C. C., Stack, G., Jiao, X., Lara-Tejero, M. and Galán, J. E. (2019) Alternate subunit assembly diversifies the function of a bacterial toxin. Nat Commun 10, 3684

52. Connell, T. D. (2007) Cholera toxin, LT-I, LT-IIa and LT-IIb: the critical role of ganglioside binding in immunomodulation by type I and type II heat-labile enterotoxins. Expert Rev Vaccines 6, 821–834

53. Scheutz, F., Teel, L. D., Beutin, L., Piérard, D., Buvens, G., Karch, H., Mellmann, A., Caprioli, A., Tozzoli, R., Morabito, S., Strockbine, N. A., Melton-Celsa, A. R., Sanchez, M., Persson, S. and O’Brien, A. D. (2012) Multicenter evaluation of a sequence-based protocol for subtyping Shiga toxins and standardizing Stx nomenclature. J Clin Microbiol 50, 2951–2963

54. Littler, D. R., Ang, S. Y., Moriel, D. G., Kocan, M., Kleifeld, O., Johnson, M. D., Tran, M. T., Paton, A. W., Paton, J. C., Summers, R. J., Schembri, M. A., Rossjohn, J. and Beddoe, T. (2017) Structure-function analyses of a pertussis-like toxin from pathogenic Escherichia coli reveal a distinct mechanism of inhibition of trimeric G-proteins. J Biol Chem 292, 15143–15158

